# Effects of reward history on decision-making and movement vigor

**DOI:** 10.1101/2021.07.22.453376

**Authors:** Shruthi Sukumar, Reza Shadmehr, Alaa A. Ahmed

## Abstract

During foraging, animals explore a site and harvest reward, and then abandon that site and travel to the next opportunity. One aspect of this behavior involves decision-making, while the other involves movement control. We recently proposed that control of decision-making and movements may be linked via a desire to maximize a single normative utility: the sum of all rewards acquired, minus all efforts expended, divided by time. If this is the case, then the history of rewards, and not just its immediate availability, should dictate how long one decides to stay and harvest reward, and how fast one travels to the next opportunity. We tested this theory in a series of experiments in which humans used their hand to harvest tokens at a reward patch, and then used their arm to reach toward a subsequent opportunity. Experiencing a history of poor rewards not only led people to forage longer, but they also reached slower toward the next reward site. Thus, reward history had a consistent effect on both the decision-making process regarding when to abandon a reward site, and the motor control process regarding how fast to move to the next opportunity.

## Introduction

We move more vigorously toward stimuli that promise a greater utility: the speed of saccades and reaching movements that are directed toward the rewarding stimulus tends to increase with the expected reward (Haith et al., 2012; Rigoux & Guigon, 2012; Shadmehr et al., 2016; Summerside et al., 2018; Takikawa et al., 2002; Thura et al., 2014). Similarly, as the effort cost of acquiring reward increases, the speed of walking and reaching tends to decrease (Gordon et al., 1994; Ralston, 1958; Schweighofer et al., 2015; J. Wang et al., 2021). Why does the prospect of greater reward invigorate our movements?

One possibility is that by controlling movement vigor the brain is attempting to maximize an ecologically important variable: the capture rate, defined as the reward that is expected at the successful conclusion of the act, minus the energetic expenditure required to perform that act, divided by time (Shadmehr et al., 2016). In this framework, faster movements are discouraged because they require greater energetic expenditure (Gordon et al., 1994; Ralston, 1958; Summerside et al., 2018). However, when there is reward at stake, faster movements save time, which reduces the discounted value of reward (Haith et al., 2012; Shadmehr et al., 2010). When greater reward is at stake, the time that is saved by a faster movement justifies its energetic cost, and thus invigorates it.

Because improving the capture rate aids survival and fecundity (Lemon, 1991), the capture rate was originally proposed by ecologists as a way to model decision-making during foraging (Stephens & Krebs, 1986; Werner, Earl and Hall, 1974). During foraging, animals harvest the reward available at the current site, and then abandon it and move to another opportunity (Cowie, 1977; Richardson & Verbeek, 1987). Optimal foraging theory operates on the notion that the decision of when to abandon harvest depends not only on the reward that is available immediately, but also on the history of the animal, i.e., the past actions and their consequences (Stephens & Krebs, 1986). For example, marginal value theorem, or MVT (Charnov, 1976), a method that finds foraging durations that maximize the capture rate, predicts that if in the past the animal has enjoyed rich rewards, then it should abandon the current site sooner. Indeed, studies have found a consistent effect of environment quality on behavioral parameters such harvesting duration and response latency (Cuthill et al., 1990; Niv et al., 2007; Perry et al., 2016; A. Y. Wang et al., 2013).

While studying saccadic eye movements and gaze behavior in humans, we suggested a generalized version of MVT to account for the fact that reward history appears to affect both decision-making, and motor control (Yoon et al., 2018). The essence of the generalized theorem is that actions, whether they be in the context of harvesting, or moving between sites, should be controlled by a single normative utility: the sum of rewards and efforts, divided by time (Figure 1a). This generalized theory (Shadmehr & Ahmed, 2020) predicts that in environments in which rewards are plentiful or require little effort, the subject should not only abandon the reward site sooner, but also move with greater vigor toward the next opportunity (Figure 1b). That is, the rewarding outcomes and effort expenditures of past actions should have a consistent effect on both decision-making during harvesting, and movement control during travel.

**Figure 1:**
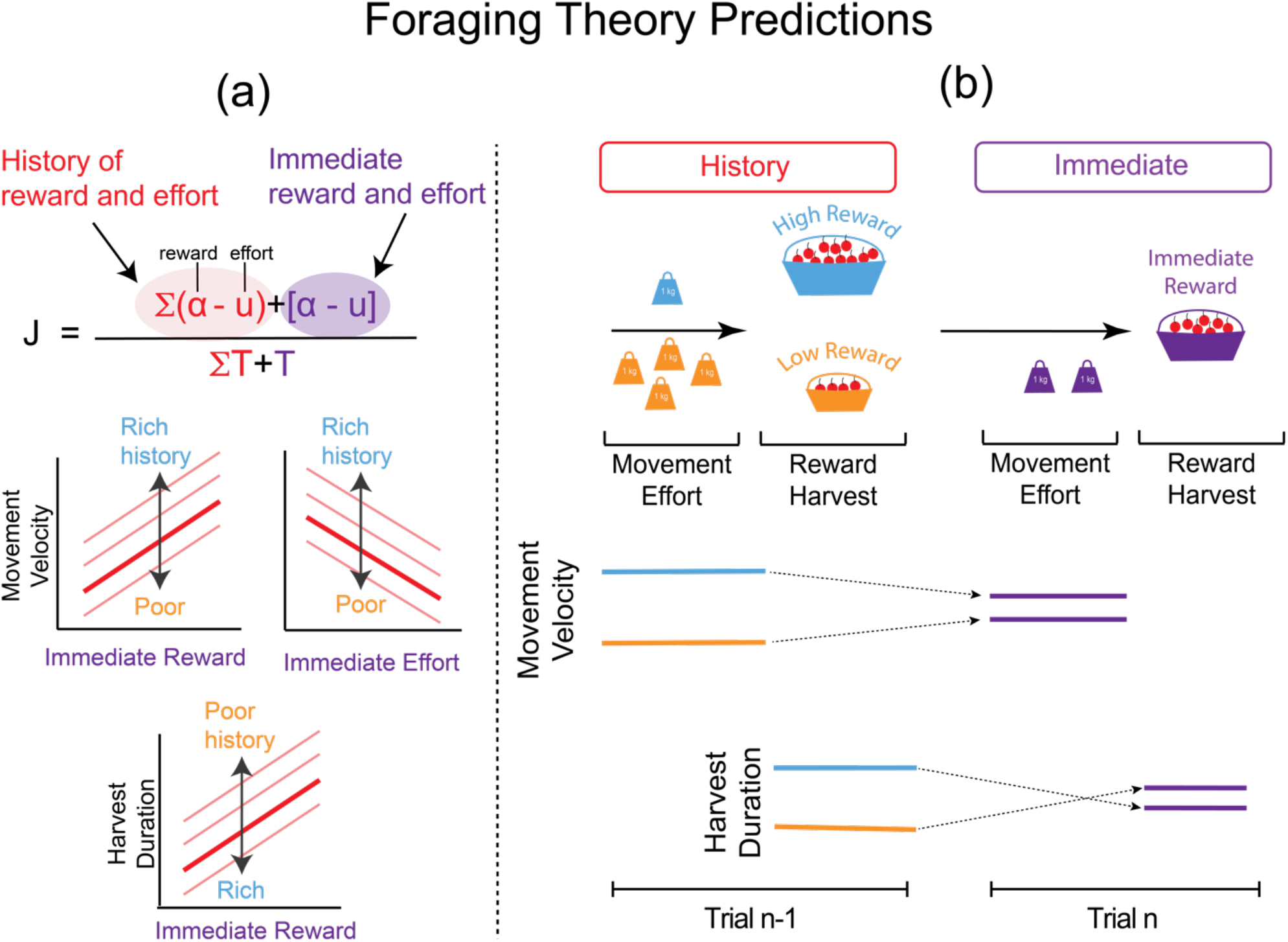
Theoretical predictions for control of movement velocity and harvest duration (a) Top: a utility, J, based on sum of prior rewards and effort costs and the immediate reward and effort costs, divided by total time. Bottom: Both immediate and history of reward and effort influence movement velocity and harvest duration. Rich environments with a history of high reward or low effort will elicit higher movement velocity, but also reduce harvest duration. (b) After a history of low effort or high reward, movement vigor toward a control immediately available reward opportunity will decrease, but the time spent harvesting that reward will increase. Note that while environmental richness is positively correlated with vigor, it is negatively correlated with harvest duration, even though an increase in the immediate reward increases both variables.

While classical MVT has been empirically tested via examination of decision-making patterns (Constantino & Daw, 2015; Cowie, 1977; Hayden et al., 2011; Krebs et al., 1974; Le Heron et al., 2020), the predictions regarding the effects of history on movement control have yet to be tested. Here, we designed foraging experiments in which we varied the history of reward and effort. Subjects used their hand to harvest tokens at a reward patch, and then used their arm to reach to the next patch. In each experiment, subjects performed these reach-and-harvest trials in various conditions: in rich environments where rewards were plentiful (or low effort expenditure associated with reaches between patches), or in poor environments where rewards were scarce (or high effort expenditure). Critically, in both environments they experienced probe trials in which we controlled both the availability of the immediate reward and the efforts required for attaining that reward. Theory predicted that in probe trials, the history of past actions, as reflected in the richness of the environment, should affect both reach vigor, and harvest duration. Indeed, following an experience of rich rewards, people not only reached with greater vigor, but also made more impulsive decisions, abandoning the harvest opportunities sooner.

## Results

Subjects held a robotic manipulandum at a reward site and applied force pulses with their hand to harvest tokens, which were exchanged for a monetary bonus at the end of the experiment. In Experiment 1 (Fig. 2a-c), there were three types of reward sites, indicated by their color, with differing reward rates (low, medium, and high, Figure 2d). As they stayed within a site and continued to harvest, the number of berries that we delivered with each pulse decayed (Figure 2d,), thus encouraging them to leave. Their decision to leave was aided by the fact that as they harvested, the location and reward rate of the next patch was available, making it possible for them to leave the current patch and reach to the next patch when they wished to.

**Figure 2:**
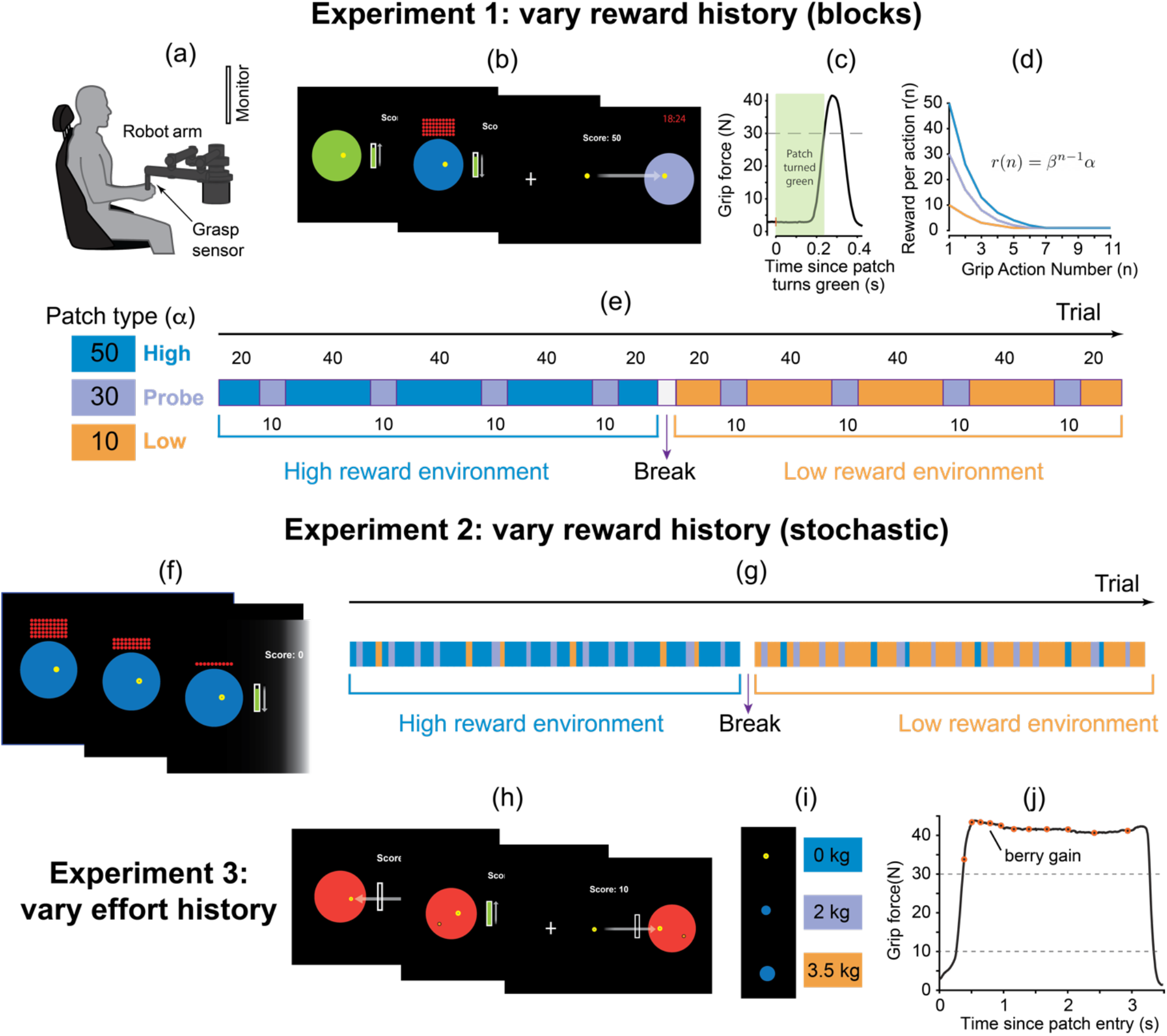
Experiment design. (a) Participants were seated in front of a computer monitor while grasping the handle of a robotic arm. The robot handle housed a sensor that measured grip force. (b) They used the robot to move the cursor and place it inside the target (“patch”) to earn reward (berries). In **Experiment 1**, we varied the amount of reward at each patch. Once in the patch, harvesting would commence by waiting for 1 second (green light), and then producing a force pulse. Each subsequent grip pulse had to wait for 1 sec and was rewarded with an exponentially reduced number of berries. The number of berries collected was added to a running score, which was displayed on the screen. Subjects were free to stop harvesting and move to the next patch at any time. The richness of each patch was cued by a specific color. (c) Example of a grip pulse resulting in berry harvest. (d) Different reward functions for the three patch types plotted with respect to the number of force pulses applied within the patch. (e) Protocol for Experiments 1 and 3. Each rectangle represents a contiguous block of trials with corresponding number of trials indicated. Color indicated trial type. Note that for experiment 1 each environment included probe trials and was conducted for a fixed amount of time (20 minutes), with the remaining time displayed in the upper right corner of the screen. In **Experiment 2** patch color did not reveal underlying reward value and patches were distributed stochastically. (f) Patch presentation stimuli for Experiment 2. (g) Protocol for Experiment 2 in which subjects experienced the three patches in random order (patch color used for protocol visualization purposes alone). In **Experiment 3**, we varied the effort required to travel between patches. (h) A red ‘patch’ served as the cue, instructing the subject to move the cursor to collect reward. Once in the patch they increased grip force to a threshold (bar height) to commence harvesting. During harvesting, grip force had to be maintained above threshold (*F*_*g*_ = 30*N*) to continue berry collection. Subjects could move out at any point to the next patch. (i) To modulate the effort cost of travel between patches, we added a mass to the reaching movements, indicated by a circle that was proportionally sized to the mass and appeared over the cursor. (j) Example grip force profile; subjects had to hold force above (*F*_*g*_ = 30*N*) to collect berries indicated by orange circles.

The key variables were harvest duration and reach velocity. We hypothesized that these variables depended not only on the immediate reward opportunities, but also their history. To control reward history, the patches were organized so that the subjects initially experienced either a poor (e.g., most patches containing a small amount of reward, Figure 2e) or a rich environment. In addition, in both environments subjects also encountered probe trials in which the immediate reward rate was kept constant (medium). These probe patches allowed us to measure the effect of past reward rate while controlling for the immediate reward. Subjects were given a fixed amount of time (20 min) to complete each environment and were familiarized with the various patches before the main experiment.

### History of high reward led to shorter harvest, and faster movements

Subjects reached faster toward patches that promised a greater reward, as shown for a representative subject in Figure 3a, and across subjects in Figure 3b (*F*(1,1104) = 80.225, *p* < 2 ∗ 10^−16^). However, when the promise of reward was equalized (probe trials), reach speed depended on reward history: following a history of high rewards, reach speed in probe trials was greater than following a history of poor rewards [peak velocity in probe trials following a high reward environment as compared to the low reward environment, *F*(1,264) = 13.661, *p* = 0.0002] (Figures 3b and 3c). Thus, reach velocity was higher in probe trials that were preceded by a sequence of high reward experiences.

**Figure 3:**
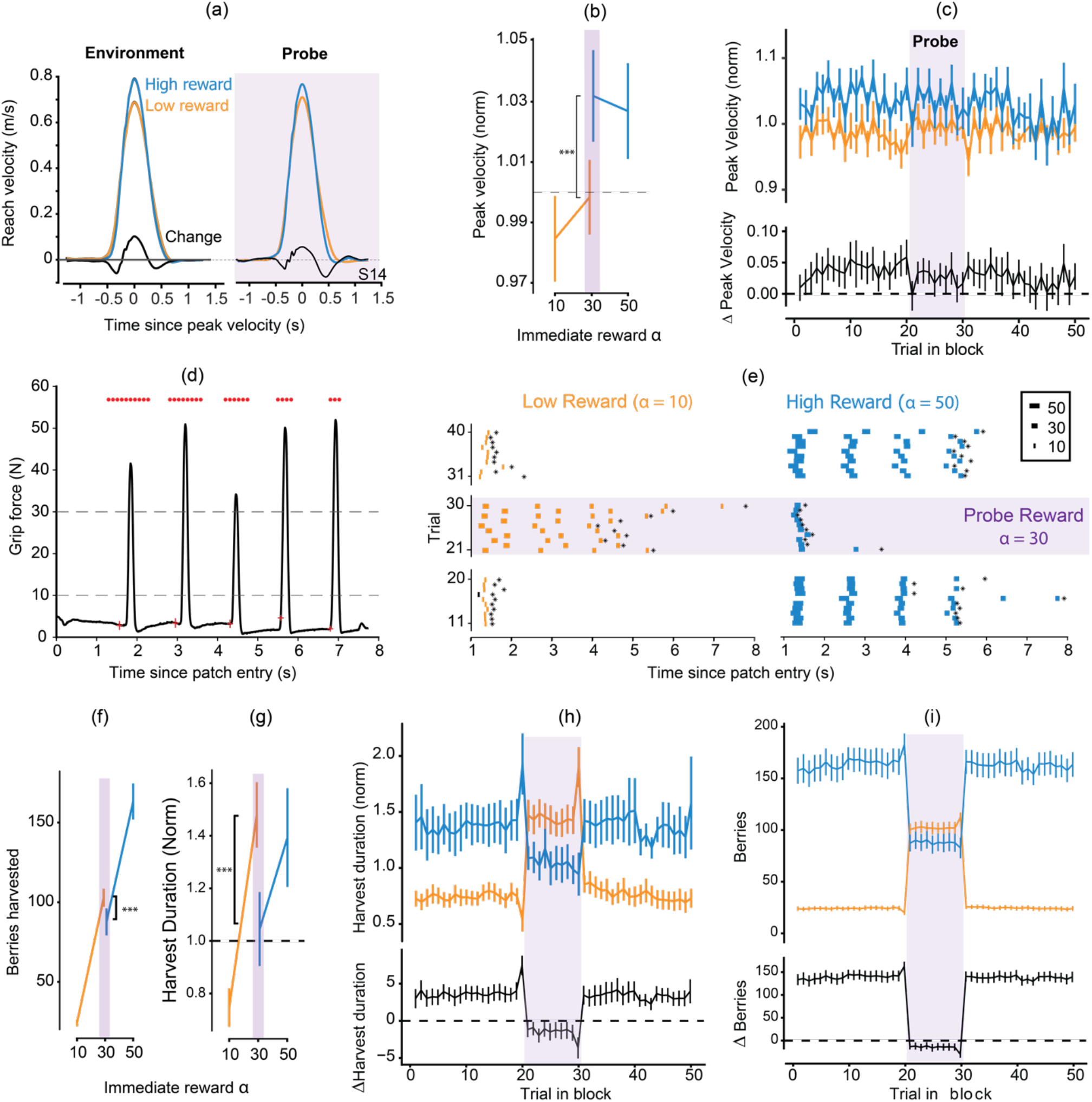
(a) Velocity profile averaged across trials belonging to each environment for a representative subject; probe trials are separated out for each environment. (b) Normalized peak velocity plotted with respect to reward in patch on the current trial; each point represents the average of the mean normalized peak velocity across all subjects. Subjects are faster in probe trials belonging to high reward environment (* p<0.05, *** p<0.001). (c) Mean peak velocity (normalized to subject average) within a block of 50 trials, averaged across subjects, plotted as a function of trial for the low (blue) and high (yellow) effort environments. Corresponding difference Δ*PV* is plotted with respect to trial number within block right below. All error bars represent standard error of the mean. (d) A sample grip force profile for an example subject #14 on a randomly selected trial to indicate the temporal course when inside a patch. In this case, subject is harvesting from a low reward patch; initial patch reward is 10 and successively decreases with grip actions. (e) Raster plot showing berries harvested in 30 representative trials per environment including for Subject #14. Each row represents a trial. Annotated trials are probe trials with equivalent reward across environments. Each point represents a harvest with the thickness of the point represents the number of berries harvested. Time of departure from patch is indicated by the asterisk for corresponding trial. (f,g) Berries harvested (w) and harvest duration (normalized to subject average; f) plotted as a function of patch initial reward *α*. Comparing metrics between the probe patches’ average between the two environments showed subjects left patches sooner after history of high reward. (h,i) Mean number of berries (g) and mean harvest duration, normalized to subject average (h) within a block of 50 trials, averaged across subjects, plotted as a function of trial for the low (blue) and high (yellow) effort environments. Corresponding difference is plotted with respect to trial number within block right below. All error bars represent standard error of the mean.

Once the hand entered a reward patch, the subjects produced a series of force pulses to collect berries (one pulse per set of berries). Figure 3d provides an example of the force pulses, and the number of berries collected following each pulse (red dots atop each pulse). The raster in Figure 3e provides an example of harvest behavior in the rich and poor environments, as well as in the probe trials (same subject as in Figure 3a). As expected, the subject stayed longer in the high reward site as compared to the low reward site, a pattern that was consistent across subjects (harvest duration was longer in the high reward patch, Figure 3g; *F*(1,1104) = 632.4, *p* < 2 ∗ 10^−16^ and more berries were collected; *F*(1,1104) = 10838, *p* < 2 ∗ 10^−16^). However, they abandoned the harvest sooner if the past was one of high rewards. Indeed, after a history of high reward the subjects chose to stay for a shorter period in the probe patch (Figure 3h, *F*(1,264) = 37.134, *p* < 3.8 ∗ 10^−9^), and collected fewer berries (Figure 3i, *F*(1,264) = 63.077, *p* < 5 ∗ 10^−17^), as compared to the probe patch that followed low rewards. In other words, given the same amount of immediate reward (i.e., probe trials), if the history was one of high reward, then the subjects abandoned the current patch earlier, thus leaving with fewer berries.

As they harvested rewards, subjects could see the location of the next patch and whether it contained high, medium, or low rewards. This allowed us to examine whether the prospect of future rewards affected behavior at the current patch. We noticed a curious change in behavior in the trial just before the onset of the probe patch: during the final harvest in the high reward patch, subjects lingered longer than normal if the next patch was a probe trial (harvest duration, trial 19 vs. 20, *t*(13) = 3.258, *p* = 0.0062). In other words, just before moving to a relatively poor patch (probe trial), subjects stayed longer than usual in the high reward patch. This transition from high reward patches to a probe patch also produced a relatively slow reaching movement (as compared to reaches to probes that were preceded by other probes, *t*(13) = −2.188, *p* = 0.0475). Similarly, when the next patch contained a relatively greater amount of reward than the current one, harvest duration was shorter than normal (final harvest in the low reward environment when the subsequent patch was probe, *t*(13) = −3.53, *p* = 0.0037). Thus, harvest duration was longer than usual if the promise of future reward was lower than the current rate.

In summary, immediate reward affected harvest duration and reach vigor: in the high reward environment people lingered longer at each patch (as compared to low reward), and then reached with greater vigor toward the next high reward patch. In addition, the history of reward affected these variables: following a low reward history, people lingered longer at the probe site, and then reached with reduced velocity to their next opportunity. Thus, as theory had predicted, a history of high reward encouraged earlier abandonment of the current reward patch, and faster movements toward the next patch.

### History of reward modulated vigor and decision-making even in the absence of explicit cues

In Experiment 1, color cues provided explicit information regarding the quality of the reward sites. Would the decision-making and movement patterns change if we were to remove these explicit cues? In Experiment 2, the subjects experienced three patch types—low, intermediate, and high reward (same patches in experiment 1), but this time in a random order (Figure 2g). However, the value of the upcoming patch was not cued; all the patches were colored blue (Figure 2f). Thus, the subjects were not informed of the environment quality explicitly and had to rely on experience. Subjects foraged in two environments, a low reward environment with 70% low reward patches, 20% medium patches and 10% high reward patches, and a high reward environment with 70% high, 20% medium and 10% low reward patches.

We found that the subjects harvested for a longer duration in the low reward environment as compared to the high reward environment (Figure 4a; *β* = −0.069; *p* = 0.0106). Critically, they spent a longer period in the intermediate probe patch when that patch was encountered in the low reward environment as compared to the high reward environment (Figure 4a).

**Figure 4:**
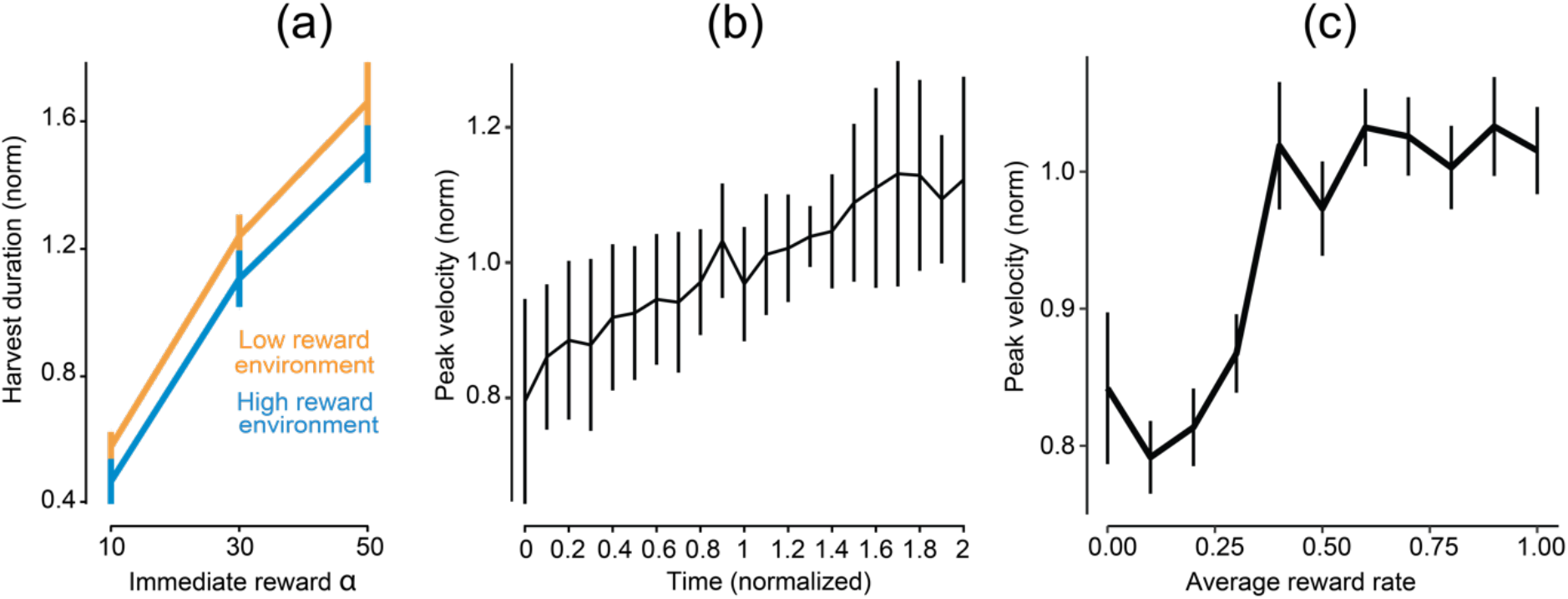
(a) Harvest duration normalized to within subject average, as a function of the reward value of the patch in the two environments. (b) Peak velocity as a function of time spent in the experiment. Time is a variable that goes from 0–2, from the start of the first environment to the end of the second. (c) Peak velocity averaged across subjects and normalized to subject average as a function of reward rate history (scaled and binned).

Focusing on vigor of reaching movements, we uncovered a confounding effect: as the experiment progressed, the subjects increased their peak reach velocity (Figure 4b). Thus, in order to determine if there was an effect of reward history on reach vigor, we first had to take into account the confounding effects of time passage. To do so, we built a linear model that for each subject considered reach vigor as a function of reward history, time, and their interactions. Once the confounding factors were accounted for, we found that peak velocity increased significantly as a function of reward history (Figure 4c; *β* = 0.0010; *p* = 0.0141).

Putting the results of Experiments 1 and 2 together, we found that reward history affected both the decision regarding how long to stay in a patch, and the vigor of movements from one patch to another. Specifically, a higher rate of reward in the environment accompanied higher vigor and shorter harvest durations.

### A history of high effort expenditure encouraged slower reaching movements

In experiment 3, we shifted our focus from history of reward to history of effort expenditure. We did this by simulating an added mass on the hand as the subjects reached between patches (via an acceleration-dependent resistive force field). Like Experiment 1, subjects foraged for discrete rewards by reaching towards them (Figure 2h). Unlike Experiment 1, once at a patch they applied a constant grip force to obtain reward (Figure 2j). As before, they were free to stop the harvest and leave for the next patch at any time (Figure 2h, third panel). They experienced three added mass conditions over the course of the experiment which were cued through the appearance of the cursor (Figures 2i). In addition, they foraged in two environments: a high effort environment (3.5 kg mass) and a low effort environment (0 kg mass). In each environment, there were intermittent probe trials (2 kg), which allowed us to compare behavior between environments when the immediate effort requirements were the same, but history of effort differed. Like Experiment 1, the trial types, in this case high, low, and medium effort, were blocked and cued (Figure 2e).

As expected, subjects reached faster when carrying a small mass as compared to a large mass (effect of environment effort on peak velocity, as illustrated by data from a single subject in Figure 5a, and group data in Figure 4b [*F*(1,1420) = 3828.4, *p* < 2 ∗ 10^−16^]). More importantly, the history of effort affected their reaching movements: reach velocity was higher in probe trials when those trials were embedded in an environment in which effort costs were low (Figure 5b, [*F*(1,1420) = 27.31, *p* < 3.03 ∗ 10^−7^]). Thus, as expected the subjects reached slower if the mass on the probe trial was greater than the trials that immediately preceded. However, this reach speed was faster if the probe trial was embedded in a low effort environment as compared to a high effort environment.

**Figure 5:**
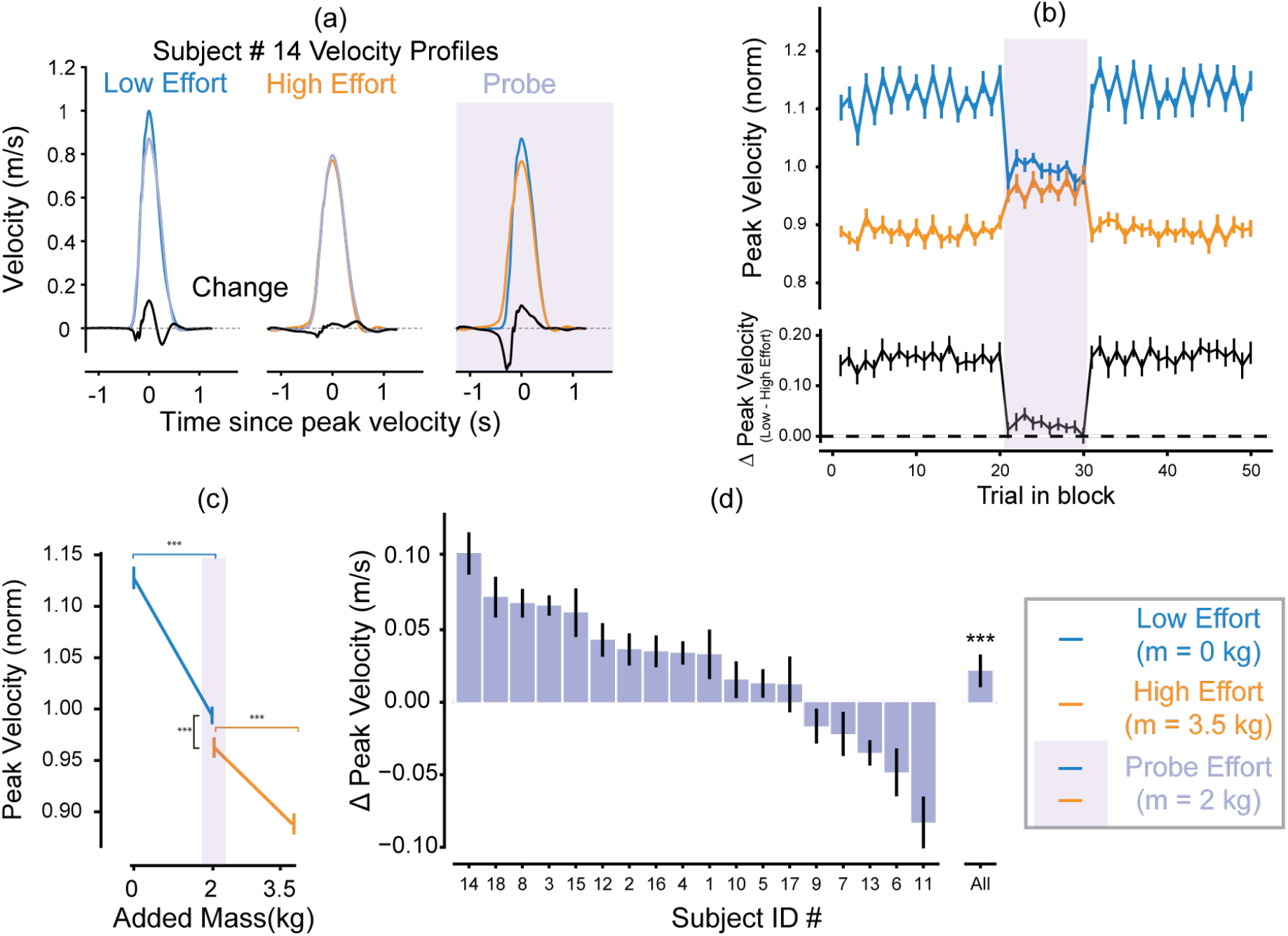
Vigor was influenced by history of effort. (a) Averaged velocity profiles for different mass conditions, 0kg (blue), 2kg (purple background) and 3.5 kg (yellow), for an exemplar subject 14. (b) Mean peak velocity (normalized to subject average) within a block of 50 trials, averaged across subjects, plotted as a function of trial for the low (blue) and high (yellow) effort environments. Corresponding difference Δ*PV* is plotted with respect to trial number within block right below. All error bars represent standard error of the mean. (c) Peak velocity plotted as a function of added mass averaged across subjects. (d) Average difference in peak velocity between corresponding probe trials in low effort environment and high effort environment 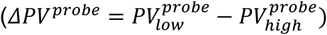 for each subject; bars are arranged in descending order of Δ*PV*. Most subjects have a mean positive Δ*PV* indicating that they move on average faster in the probe trials belonging to the low effort environment.

Interestingly, the effect of effort history on vigor was not uniform across all subjects (Figure 5d). Specifically, five of the 18 subjects presented a slight or strong increase in probe trial vigor in the high effort environment. One possible explanation for this is that, despite the available reward and reward rate being uniform in patches across both environments, subjects valued the reward in the high effort environment more by the virtue of having spent more effort to acquire it. This is addressed further in the Discussion section.

Upon entering a patch, subjects increased their grip force to the threshold required for harvesting. This threshold remained constant despite the varying effort requirements (i.e., masses) of the environment. Interestingly, subjects altered the rate at which they increased their grip force (grip ramp-up phase in Figure 6a) based on the effort history of the environment: in the probe trials that were placed in the high effort environment, the rate of increase in grip force tended to be slower.

**Figure 6:**
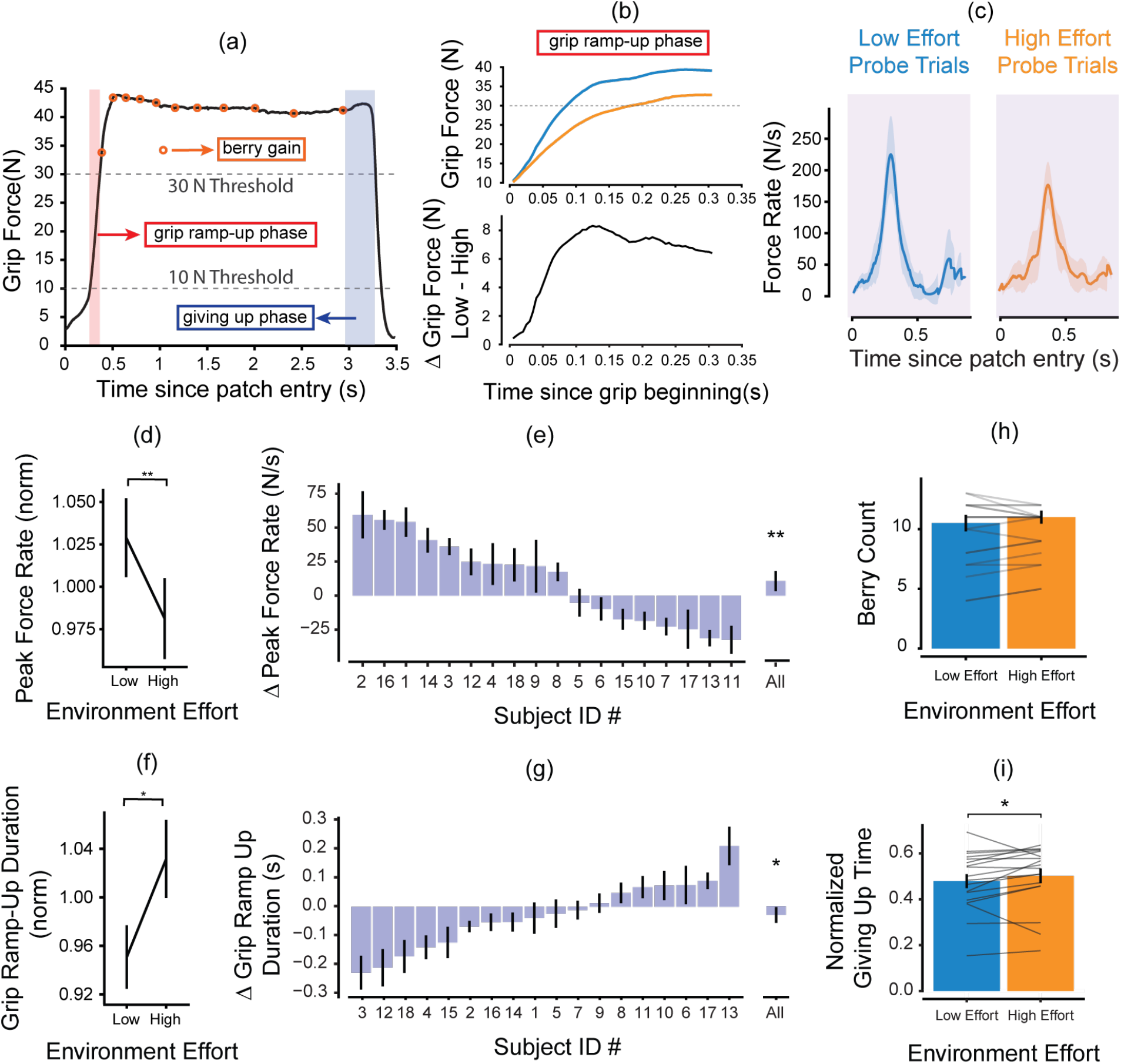
Harvest behavior is modulated by history of effort (a) A sample grip force profile for an example subject 14 on a randomly selected trial to indicate the temporal decision phases when inside a patch. (b) Zooming in on the grip ramp up phase we compare the grip force profiles between two probe trials for subject 14, one from the low effort environment (blue) and the other from the high effort environment (orange). (c) Force rate profiles for subject # averaged across probe trials within the two environments; peak force rate in the probe trials belonging to the low effort environment was higher than that of the high effort environment. (d) Peak force rate – the maximum rate of force generation during ramp-up phase– normalized to subject average, was lower in probe trials belonging to the high effort environment. (e) Most subjects had a positive ΔPeak force rate. (f) Grip Ramp-up Duration – which is computed as the duration from when the force is at 10N to when the force is at 30 N – increased in the probe trials belonging to the high effort environment as opposed to the low effort environment. (g) Mean difference in harvest reaction time in probe trials between environments. As expected, ΔHarvest reaction time is negative, for most subjects. (h) Berry count modulation by environment was not statistically significant. (i) Giving-up time, normalized to the wait time between the last two berries, increased in the high effort environment.

Two examples of harvesting trials are shown in Figure 6b. This subject increased their grip force faster in the context of the low effort environment. We quantified this behavior by measuring the rate of force increase as well as the duration of time it took to ramp force from 10 N to the threshold of 30 N. (Subjects were required to maintain their force below 10 N before they enter the patch thereby determining the lower threshold.) We found that on average, peak rate of force was higher in trials belonging to the low effort environment as opposed to the high effort environment [example subject Figure 6c; two-way repeated measures ANOVA, main effect of environment on peak force rate *F*(1,264) = 5.714, *p* = 0.0175]. When we focused on probe trials, we found that peak force rate in the context of the low effort environment was higher than in the context of the high effort environment [Figure 6d–e; two-way repeated measures ANOVA, effect of environment effort on probe trials, *F*(1,1382) = 17.442, *p* < 0.001]. When considering the correlated metric of grip ramp-up duration we see an interaction between environment effort and trial type [two-way repeated measures ANOVA on all trials reveals a slightly significant interaction between environment effort and trial type, *F*(1,264) = 3.851, *p* = 0.05076]. Focusing once again on just the probe trials we found a significant effect of environment effort on grip ramp-up duration [Figures 6f–g; *F*(1,1382) = 8.909, *p* = 0.00289]. From Figures 5d and 6e, we notice that there is a significant overlap among subjects who had a higher peak force rate in the high effort environment and those who moved faster in the high effort environment. Specifically, subjects 6, 7, 11 and 13 who have higher peak velocity in the high effort environment also grip faster in the high effort environment. This provides more evidence in support of the notion that these subjects view the high effort environment as better than the low effort environment.

Following the ramp-up phase, subjects harvested berries. In contrast to the grip ramp-up phase, we found no consistent effect of environment effort (*p* = 0.690) on number of berries collected (Figure 6h), even when focusing on only the probe trials (*p* = 0.621). Finally, we focused on the behavior as the subjects ended the harvest period and reduced their grip force, termed giving-up phase (Figure 6a), or the duration that subjects waited in a patch gripping after collecting their last berry. Since the duration between consecutive berries increased with time spent in the patch, this metric was normalized to the time experienced waiting between the second-to-last and last berries. We found that this normalized giving-up time was modulated significantly by environment effort (Figure 6i [two-way repeated measures ANOVA, main effect of effort; *F*(1,264) = 5.437, *p* = 0.0205]). That is, in the high effort environment, the subjects waited longer before giving up. Importantly, we found no significant effect of environment [*F*(1,1382) = 2.088, *p* = 0.149].

In summary, for identical immediate effort and reward opportunities, movement vigor tended to be modulated by the subject’s past experience: following a history of high effort expenditure, people reached with reduced velocity in probe trials. Following a history of high effort, subjects did not significantly modulate the amount of reward that they harvested. However, they took longer to begin harvesting reward (grip ramp-up period) and waited longer before ending the harvest and moving on to the next patch (giving-up phase).

## Discussion

During foraging, the decision-making and movements of an individual may be linked via a single normative utility: the sum of rewards acquired, minus efforts expended, divided by time (Shadmehr & Ahmed, 2020). A prediction of this theory is that the history of reward should affect decision-making during harvesting, and movements during travel. Indeed, we found that people harvested longer and moved slower following a history of low rewards (or high effort), and harvested for a shorter duration and moved faster following a history of high rewards (or low effort). History of effort had a similar effect on reach vigor: following a history of high effort expenditure, people reduced their reach velocity in probe trials. However, unlike the theoretical predictions, history of high effort did not encourage longer harvest durations. Rather, it slowed the force production patterns that initiated and ended the harvest.

History of reward and effort influenced both the choice of how long to harvest, and the control of movement vigor to the next opportunity. There is a large body of work demonstrating that humans and other animals modulate their harvest duration in patches based on reward history in accordance with MVT (Constantino & Daw, 2015; Hills et al., 2012; Krebs et al., 1974; Le Heron et al., 2020; Wikenheiser et al., 2013; Wolfe, 2017; Yoon et al., 2018). In contrast, the influence of reward history on movement vigor is considerably less understood. Our observation here further link the control of movements and decision making.

In the context of immediate reward, recent work has revealed a link between movement vigor and preference. For instance, it has been shown that humans (Sackaloo et al., 2015; Summerside et al., 2018) and other animals (Mosberger et al., 2016; Opris et al., 2011) are willing to be energetically inefficient by reaching faster and even reacting sooner towards increased reward. This invigoration has also been noted in the increased speed (Haith et al., 2012; Reppert et al., 2015) and shorter reaction times (Kawagoe et al., 1998; Takikawa et al., 2002) of saccades. Added effort also appears to slow down reaching and walking movements (Gordon et al., 1994; Ralston, 1958; Shadmehr et al., 2016). Moreover, when choosing between options immediately available, movement vigor reveals our underlying subjective valuation. People saccade faster to the option that they value more (Korbisch et al., 2019; Reppert et al., 2015; Yoon et al., 2018). Our contribution in this work is to demonstrate that this connection between movement vigor and decision making is observed not only for immediate reward, but for the history of reward as well. Specifically, we observe a consistent effect of local availability of reward as well as the history of reward on the control of vigor.

In the field of motor control, the history of actions performed has also been shown to play a role in determining the kinematics of subsequent movements. For example, movement repetition leads to an experience-dependent learning process whereby the brain learns to reduce variability of performance towards previously repeated targets (Diedrichsen et al., 2010; Verstynen & Sabes, 2011). This has been postulated to be a trial-by-trial learning process of the statistics of the previous actions with a bias-variance tradeoff strategy explaining selected action kinematics in relation to allowed preparation time. This bias has been observed in the speed as well as reaction times of repeated movements (Hammerbeck et al., 2014; Mawase et al., 2018). This use-dependent effect is however separable from another history dependent phenomenon observed whereby subjects use a cognitive, predictive strategy when deciding subsequent movements in the presence of uncertainty (Marinovic et al., 2017). But what about the effect of dissimilar actions and their outcomes on subsequent movement kinematics? The optimal foraging framework thus seeks to explain durations of different actions, by defining a unifying ecological utility that the actions and their durations modulate. Our findings demonstrate that both the local context of reward and effort as well as the history of reward and effort significantly affect vigor.

Consistent with the predictions of the generalized MVT, people traveled more slowly following a history of high effort. These results are however at odds with those presented by Yoon et al. (Yoon et al., 2018) for saccade vigor. They found that after a history of high effort, imposed by high eccentricity of images, while gaze increased (increase harvest duration), saccade velocity also increased. In other words, subjects made faster saccades in a poor environment when moving across equivalent distances between images of identical eccentricity. This result was discussed by the authors as being more aligned with a framework in which effort expenditure elevated the subjective reward value in the environment. This elevation of the reward due to effort expenditure is referred to as *justification of effort* whereby the same reward higher is valued more if more effort was required to obtain it (Klein et al., 2005; Tricia, S et al., 2000). Though Yoon et al.’s (Yoon et al., 2018) results contrast with ours. In Experiment 3, we found that some subjects moved faster in the high effort environment as opposed to the low effort environment in the probe trials. The justification of effort phenomenon could explain these inter-individual differences.

In the context of modulated travel effort, we found that some aspects of harvest behavior were modulated by effort history (Figure 5). Specifically, we also find modulation in “giving-up” duration, a metric that has been used to define how long an animal remains in a patch without reward before leaving. In previous discrete reward foraging scenarios, an increase in giving-up duration has been seen to reflect the forager’s perception of environment quality or capture rate. For instance, birds have been known to have a longer giving-up time in poorer environments (Krebs et al., 1974). As in the literature, and per the prediction of MVT, we find that subjects wait longer before giving-up in the high effort environment.

The neural correlates of reward history remain poorly understood. Dopamine levels in the nucleus accumbens correlate with reward rate, as well as shorter response latencies (Mohebi et al., 2019). However, the source of the increase remains elusive since in that same task, tonic firing rate of dopaminergic neurons do not track the history or reward or punishment (Cohen et al., 2015; Mohebi et al., 2019). A clue to this puzzle may lie in another neurotransmitter: serotonin. Tonic firing rates of serotonergic neurons can reflect history of reward, with other serotonin neurons encoding history of punishment (Cohen et al., 2015). Artificial activation of serotonin neurons leads to increased harvest durations in a foraging task (Lottem et al., 2018), as well as reduced movement vigor (Correia et al., 2017; Seo et al., 2019). Taken together, these results suggest that serotonin may play a role in linking decision making and movement vigor during foraging via an encoding of reward history.

Not all of our findings conformed with our theory. Previous work has found that after longer travel delays, birds will stay longer and collect more food in subsequent patches with depleting rewards (Cuthill et al., 1990). This has been shown in humans as well, where individuals harvesting reward from virtual trees will collect more apples after a longer travel delay (Constantino & Daw, 2015). These imposed travel delays seek to reduce the average capture rate of the foraging environment and therefore led to longer stay times. However, in our study we see a lack of modulation of the absolute number of berries collected in response to changing environment quality due to change in travel effort. In our protocol, each berry is associated with a short high-pitched beep, providing a salient signal regarding the number of berries harvested. We believe that subjects settle on several berries that leads to a predictable auditory pattern, that in turn acts as a cue as to when to leave a patch. In other words, subjects learn to expect a certain number of berries irrespective of reward rate due to the auditory feedback, rather than deciding on a per-patch basis. There has been some conflicting evidence for (Mcnair, 1982) and against (Krebs et al., 1974) the phenomenon of predators expecting a set number of prey, or hunting by expectation as opposed to a strategy based on MVT. Nevertheless, in Experiment 3, the increased saliency associated with the number of berries may prime subjects to collect a fixed number between environments. Notably, in Experiment 1, where auditory feedback was not linked to number of berries, we observed a consistent effect of environment on berries harvested, in accordance with the theoretical predictions.

The MVT framework presents an implicit, circular solution (Stephens & Krebs, 1986) for the optimal durations it prescribes, which are dependent on the very quantity it tries to optimize. This leads to the assumption that the forager has complete knowledge of past and future rewards. Therefore, the theorem is unable to account for transients where the agent needs to learn the quality of the environment as a parameter that can be updated (Mcnamara & Houston, 1985). In our data, we see that the environment’s influence of vigor appears to get washed out by the final block of the environment with the difference between the probe trials’ vigor vanishing to zero. In addition, the theorem does not allow for stochasticity in the environment, as it does not account for fluctuating beliefs of the forager regarding the quality (Oaten, 1977). This has been noted in the literature by several studies that have presented alternatives (Green, 1980; Mcnamara & Houston, 1985; Oaten, 1977; Pyke, 2019). Yet, despite these shortcomings, empirical studies show that humans and animals generally behave according to the theorem’s predictions. There is now a push to understand the underpinnings of everyday decisions from an ethological standpoint (Hayden, 2018; Mobbs et al., 2018) through more naturalistic experiment designs that better reflect decisions faced by individuals every day. Our attempt to integrate motor control into this theory may provide a useful way to link decision making with movement control.

In conclusion, for identical immediate reward and effort opportunities, harvest duration and movement vigor were modulated by the history of reward and effort. People harvested longer and moved slower following a history of low reward, and conversely, harvested for a shorter duration and moved faster following a history of high reward. In accordance with the maximization of a normative utility, history of reward and effort exerted a consistent effect on not only the choice of how long to stay in the current patch but how fast to move to the next one.

## Data Availability Statement

All collected data and corresponding code for statistical analysis are available at https://github.com/ssukumar/foraging_code.

## Materials and Methods

### Experiment Protocol and Design

#### Subjects

Forty-four subjects (age= 24.25 ± 3.5 years, 19 female) participated in the study, fourteen in experiment 1 and ten in experiment 2 and twenty in experiment 3. All subjects were healthy with no recent injuries or known pathologies. Consent was obtained in accordance with the University of Colorado Institutional Review Board. Subjects were paid a base amount of $10 an hour, but their final compensation depended on their performance. Two subjects were unable to complete the experiment due to their selection of very long harvest duration in experiment 3 and therefore results for this session presented are from the 18 subjects who completed all the trials in the experiment.

#### Apparatus and Data Acquisition

To test predictions of MVT in humans, a foraging task involving arm reaching movements was designed. Subjects were seated in a chair with full back support and made horizontal, planar reaches while grasping the handle of a robotic manipulandum (InMotion 2; Interactive Motion Technologies, Shoulder-elbow robot 2) as seen in Figure 2a. By moving the robot handle subjects controlled the movement of a cursor on the monitor placed at eye-level. The monitor displayed a game screen in which they were cued to move to different targets, causing them to make reaches in different directions in the horizontal plane. The end of the robot handle was attached with a grasp sensor that measured force with which subjects gripped the handle. Additionally, the robot could produce forces; here we leveraged this by having the robot produce acceleration-dependent resistive forces to simulate the effect of adding mass to the reach in the horizontal plane for experiment 2 (Equation 1). Robot forces were inactive during experiment 1.

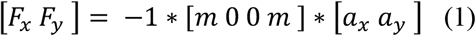

Reach position, velocity, acceleration, and corresponding grip force of every trial was recorded by the robot at a frequency of 200 Hz. The force field produced by robot motors during a reach was also continuously updated, based on the current acceleration as recorded by the accelerometer, at a rate of 200 Hz.

#### Experiment 1

This experiment emulated a patch foraging task in which subjects could move between patches by making reaching movements with their dominant arm (Figure 2a). On the screen in front of them the foraging task appeared; subjects were instructed to move their cursor into a circular patch of reward (Figure 2b). Once their cursor was in the patch, they could collect reward by making pulsed grip forces: they waited for the patch to turn green and then they could produce a force pulse (minimum of 30 N), and upon return to less than 10 N, they received rewarding ‘berries’. Subjects could wait for the patch to turn green to collect more reward or move on to the next patch at any time. Their total earnings from a patch were added to a cumulative score counter at the top of the screen.

Overall, subjects experienced three patch types with differing reward, each indicated by a color cue. Reward was dispensed per grip according to an exponential decay function, jittered by gaussian noise *r*(*n*) ∼ *N*(*β*^*n* − 1^ *α*, 1), where *n* is the pulse number, *α* represents the maximum reward that can be obtained from a pulse, and *β* represents the decay rate fixed at 0.8 for the entirety of this experiment. The first pulse always resulted in the maximum reward, *r*(1) = *α* (Figure 2c, d).

Before the main experiment, subjects were familiarized with the patch types with 30 trials, 10 for each reward value. Subjects were questioned about which patch was the most rewarding to ensure they had internalized the color-reward mapping. If they responded incorrectly, they were re-familiarized. No subject required more than 2 familiarization sessions; familiarization data were not included in any analyses presented.

#### Experiment 2

In this experiment we removed the color cues that indicated the magnitude of reward available in each patch (Figure 2f). Moreover, the subjects experienced patches in a randomized order, rather than in a block (Figure 2g). In the low reward environment subjects experienced the low reward patch 70% of the time, the intermediate reward patch 20% of the time and high reward patch 10% of the time. In the high reward environment, the proportion of low and high reward patches was swapped, with the intermediate reward patches remaining at 20%. The intermediate reward patch served as probe trials.

#### Experiment 3

In this experiment, the effort of travel was modulated by adding mass to the reaches between the patches. A brief familiarization session was conducted to acquaint subjects to the passive inertial forces of the robot arm as well as the acceleration-dependent force-field. Subjects were then required to collect berries in a task similar to Experiment 1. The available patch was displayed by a red circle. Subjects moved the robot handle and placed a cursor inside the patch. Once inside the patch, they produced a grip force that was specified by an indicator next to the patch circle (Figure 2i). Harvesting began when the grip force reached the minimum required force. In contrast to the discrete force pulses required in Experiments 1 and 2, subjects were required to continuously hold their force at the required level to harvest berries and reduce it only when they intended to move out of the patch. Berry harvesting was indicated by an animation: an orange circle quickly appearing and disappearing, accompanied by a high-pitched beep. Berries were harvested at a declining rate while the grip force was maintained above the minimum level. The total number of berries collected over a duration *t*_*h*_ was specified by the function 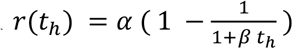.

Essentially, the time between successive berries at any given instance was calculated from the instantaneous reward rate resulting in consecutive berries having longer durations between them as time spent in the patch increased. Here, *α* represents the maximum reward and *β* represents rate of decline in reward harvested. They were told that they were free to leave the current patch at any point in time towards the cued position where a new patch would appear. Once they exited a patch, it disappeared, and a new replenished patch appeared in the new location (Figure 2h). Patches always appeared at the same two positions thereby keeping the travel distance constant at *d* = 30 *cm* across the entire experiment. Subjects (n=18) experienced the foraging task in two main blocks of trials: low mass and high mass environments. Travel effort was modulated by changing the added mass *m* to the reach period and harvest effort was represented by the amount of grip force that was to be maintained to ensure berry “consumption”.

Subjects experienced three effort levels with three different values of added mass. This was cued to the subject by means of cursor appearance (Figure 2i). The protocol entailed foraging in two environments. The low effort environment entailed no added mass (*m* = 0 *kg*) for most trials. The high effort environment entailed large added mass (*m* = 3.5 *kg*). Both environments had probe trials in which the added mass was *m* = 2 *kg*. Each environment contained two hundred trials that was divided into four sub-blocks of fifty trials each, not including 30 trials in which subjects were familiarized with the parameters of the environment. Subjects were not informed of the total number of trials in each environment. Rather, they were told that the total session duration was about one hour and fifteen minutes, including instruction and consent procedures. Within each sub-block of fifty, the middle ten trials were designated as probe thereby leading to forty total probe trials in each environment. There was no discernable break between trials in the two environments. For all trials in both environments, including probe trials, the minimum amount of grip force required to harvest berries within each patch was fixed (*F*_*g*_ ≥ 30*N*).

When travelling between the two patches, the amount of added mass was indicated by means of a modified cursor. Subjects were also instructed not to increase the grip force during travel between patches, to decouple harvest and movement efforts. If they chose to travel with increased grip force (> 10 N), the game paused, a message appeared on screen asking them to reduce their grip force. Subjects were also informed that every 10 berries collected would correspond to 1¢ in monetary bonus, to increase task engagement.

In both experiments, environment order was counterbalanced across subjects.

*Behavioral metrics of movement and harvest:* In Experiment 1, our primary question concerned the modulation of movement vigor as a function of reward history. Movements took place between patches placed on the horizontal axis, and thus we computed peak reach velocity in the horizontal direction and used it to quantify the vigor of the movement. For harvest behavior, we use two metrics. One was the number of berries harvested per patch, the other was the duration of harvest within a patch.

In Experiment 2, we removed all explicit cues that indicated reward value. Thus, we computed a post-hoc variable of average reward rate (Equation 2): 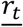 represents the average reward rate at time *t* and *R*_*t*._ represents the current patch reward. Since subjects must wait one second to harvest the number of berries *R*_*t*_, this also represents a rate. Thus, average reward rate was computed as follows:

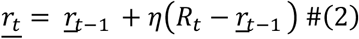

To find the value of the variable *η*, we considered the fact that the overall harvest duration was modulated by the reward rate (significant main effect of environment reward in table 2). Thus, used the dependence of harvest duration to the reward rate to find *η*. We built a linear mixed effects model (Equation 3) and used a continuous reward rate function (Equation 2) as a predictor instead of the binary variable indicating environment reward.

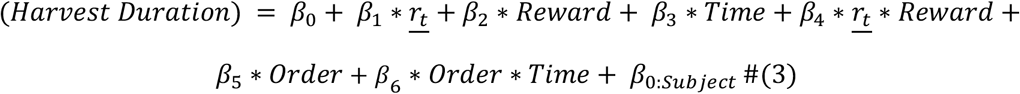

We then optimized over different values of *α* to maximize the log-likelihood of the model fit in equation 3. *η* values were restricted to be between [0,2] to maintain stability of the model with respect to the reward rate computation. The final optimal *η* was 0.036 and the corresponding model yielded a log-likelihood of 379.6 and AIC score of −741.16. Table 1 has the corresponding coefficients for harvest duration for this model; once again we show only those predictors that have a significant effect on the dependent variable.

**Table 1:**
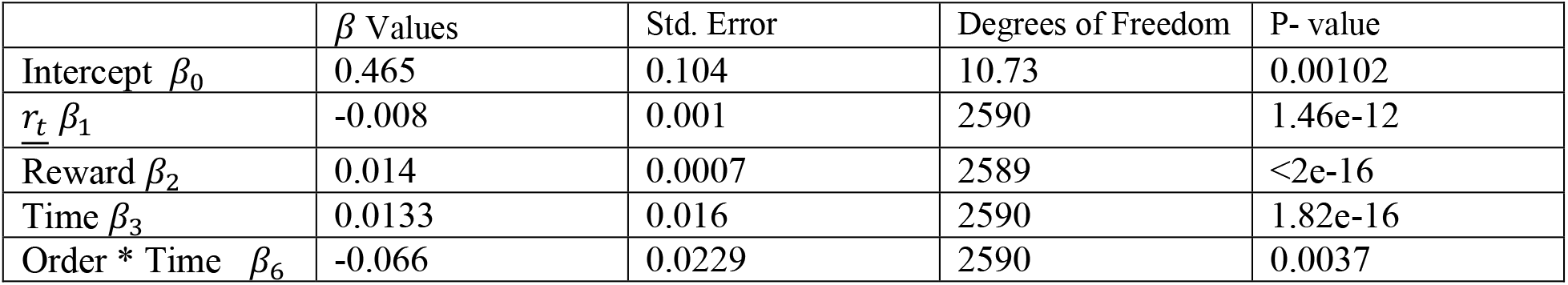
Results for model in equation 3; only significant predictors are included

| | $\beta$ Values | Std. Error | Degrees of Freedom | P- value |
| --- | --- | --- | --- | --- |
| Intercept $\beta_0$ | 0.465 | 0.104 | 10.73 | 0.00102 |
| $\underline{r}_t \beta_1$ | -0.008 | 0.001 | 2590 | 1.46e-12 |
| Reward $\beta_2$ | 0.014 | 0.0007 | 2589 | <2e-16 |
| Time $\beta_3$ | 0.0133 | 0.016 | 2590 | 1.82e-16 |
| Order * Time $\beta_6$ | -0.066 | 0.0229 | 2590 | 0.0037 |

**Table 2:** Results for model in equation 4; only significant predictors are included

| | $\beta$ Values | Std. Error | Degrees of Freedom | P- value |
| --- | --- | --- | --- | --- |
| Intercept $\beta_0$ | -0.595 | 0.143 | 10.00 | 0.0019 |
| $r_t$ $\beta_1$ | 0.0010 | 0.0004 | 2593 | 0.0141 |
| Time $\beta_3$ | 0.0870 | 0.0106 | 2593 | 2.74e-16 |
| Order * Time $\beta_5$ | 0.1177 | 0.0144 | 2593 | 4.57e-16 |

The coefficients are qualitatively like that obtained for harvest duration in the previous section; notably the optimal *η* predicts an inverse relation between harvest duration and reward rate (which has been shown in the literature for both human and animal behavior). Therefore, we are now left with a more granular measure for reward rate based on subjects’ harvest behavior. We next build a mixed effects model for peak velocity based on equation 2, but once again replacing environment reward with the reward rate metric we just computed with the optimal *η* (shown in equation 4^1^). This mixed-effects model yielded an AIC score of - 2700.4. The results for the effects are shown in table 4.

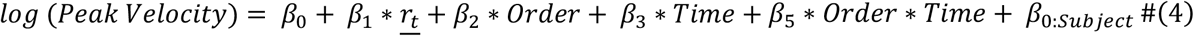

For Experiment 3, our primary hypothesis concerned how movement vigor was modulated by changing the history of effort. We quantified behavior using four metrics: harvest reaction time, peak force rate, number of berries, and giving-up time. Rate of force was computed as the numerical derivative of force over the course of the trial which was then filtered through a low-pass Butterworth filter with a 10*Hz* cut-off frequency. Since this was a grip-and-hold task. The harvest reaction time was defined as the duration from when subjects increase grip force above 10 *N* until the point at which they reach the required threshold of 30 *N*. Peak force rate was the maximum rate of force generation during the grip ramp-up period. Together, harvest reaction time and peak force rate determined the speed with which the subjects began harvest following arrival in the patch. The number of berries per patch was computed as the total number of berries in a patch that subjects collect. Finally, we computed giving-up time as the duration after the last berry was collected until they reduced the grip force below the minimum. Because berries were dispensed after increasingly long intervals, this quantity was normalized to the duration between the second to last and last berries collected.

#### Statistical Analyses

Each environment had two trial types— exemplar trials (representing the actual attributes of the environment) and probe trials (different from exemplar but equivalent across environments). Our theory predicts an effect of immediate reward and effort, i.e., if reward increases, or effort decreases, it should equivalently affect vigor as well as harvest duration. This was tested by comparing exemplar trials across both environments. The critical test comes from the comparison of probe trials between the two environments. We therefore combined all the trials across each environment and obtained average behavior across the block of fifty trials. This captures temporal behavior as well differences between environments. We performed a two-way repeated measures ANOVA on each trial type to obtain the effects of environment as well as the effect of trail number within block. The two equations representing these ANOVAs for each trial type are presented (Equations 5 & 6). *Met* represents the above metrics for movement vigor as well as harvest duration.

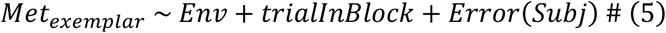

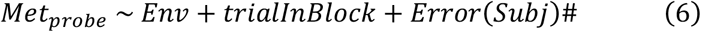

We also completed 6 post-hoc t-tests based on observing the data to test the effect of the transitioning between exemplar and probe trials for each metric. This helped us determine if the sudden changes in harvest duration and peak velocity before the transition compared to neighboring trials of the same trial type were statistically significant. For harvest duration we compared trials 29 and 30 and trials 30 and 31 in the block average for each subject in the low reward environment. Correspondingly in the high reward environment we compared trials 19 and 20 as well as trials 20 and 21. For peak velocity in the low reward environment we compared trials 30 and 31 as well as trials 31 and 32. And for the high reward environment we compared trials 20 to 21 and trials 21 to 22 within the block averages for each subject.

Additionally for Experiment 2, we ran an initial mixed effects regression to determine the effect of environment on our behavioral metrics we ran two regression models, one for harvest duration and one for peak velocity according to equations 7 and 8.

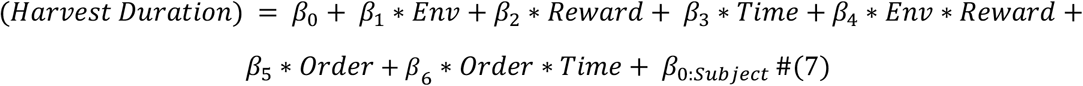

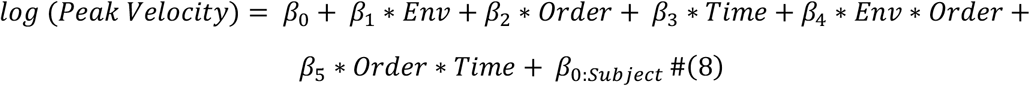

The corresponding results of the regression on harvest duration and peak velocity are available in tables 3 and 4 respectively.

**Table 3:**
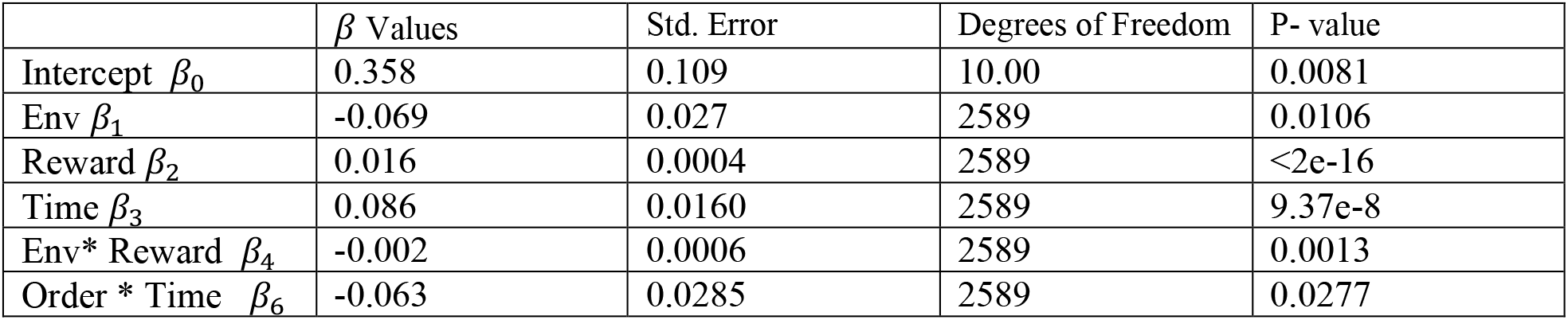
Results for model in equation 7; only significant predictors are included.

| | $\beta$ Values | Std. Error | Degrees of Freedom | P- value |
| --- | --- | --- | --- | --- |
| Intercept $\beta_0$ | 0.358 | 0.109 | 10.00 | 0.0081 |
| Env $\beta_1$ | -0.069 | 0.027 | 2589 | 0.0106 |
| Reward $\beta_2$ | 0.016 | 0.0004 | 2589 | <2e-16 |
| Time $\beta_3$ | 0.086 | 0.0160 | 2589 | 9.37e-8 |
| Env* Reward $\beta_4$ | -0.002 | 0.0006 | 2589 | 0.0013 |
| Order * Time $\beta_6$ | -0.063 | 0.0285 | 2589 | 0.0277 |

**Table 4:** Results of linear mixed model given in equation 8. Only the significant predictors are included in the table.

| | $\beta$ Values | Std. Error | Degrees of Freedom | P- value |
| --- | --- | --- | --- | --- |
| Intercept $\beta_0$ | -0.597 | 0.143 | 10.00 | 0.0019 |
| Time $\beta_3$ | 0.113 | 0.014 | 2590 | 8.90e-14 |
| Env* Order $\beta_4$ | 0.11 | 0.023 | 2590 | 1.16e-5 |
| Order * Time $\beta_5$ | 0.169 | 0.020 | 2590 | <2e-16 |

We use a statistical threshold of *α* = 0.05 for all comparisons. P-values are reported exactly unless they are less than 0.001 in which they are reported as such. All the F-statistics, confidence intervals and p-values are reported in the results for each metric.

## Footnotes

1 The equation is slightly different to the one in equation 2, i.e., there is no interaction term added between reward rate and Order due to very high collinearity between the predictor variables.

## Notes

### Competing Interest Statement

The authors have declared no competing interest.

### Summary of Updates

Additional experiment was conducted to provide further support for hypotheses.

